# Mechanistic and compositional studies of the autophagy-inducing areca nut ingredient

**DOI:** 10.1101/669648

**Authors:** Chang-Ta Chiu, Shyun-Yeu Liu, Ching-Yu Yen, Meng-Ting Tsai, Huei-Cih Chang, Young-Chau Liu, Mei-Huei Lin

**Affiliations:** Department of Dentistry, Tainan Municipal An-Nan Hospital China Medical University, Tainan, Taiwan; Oral and Maxillofacial Surgery Section, Chi Mei Medical Center, Tainan, Taiwan; Department of Dentistry, Taipei Medical University, Taipei, Taiwan; Department of Dentistry, National Defense Medical Center, Taipei, Taiwan; Department of Biotechnology, Chia Nan University of Pharmacy, Tainan, Taiwan; Department of Medical Research, Chi Mei Medical Center, Tainan, Taiwan; Division of Natural Science, College of Liberal Education, Shu-Te University, Yanchao District, Kaohsiung City, Taiwan, Republic of China

**Keywords:** areca nut, autophagy, Atg5, shRNA, caveolin, endocytosis, proteasome

## Abstract

Areca nut (AN) is a popular chewing carcinogen worldwide causing a variety of diseases such as oral and esophageal carcinomas. We previously found that the partially purified 30-100 kDa fraction of AN extract (ANE 30-100K) induces autophagy in oral carcinoma OECM-1 cells and some other different types of cells. Since autophagy is known to play important roles in tumor establishment and development, the underlying mechanisms of ANE 30-100K-induced autophagy (AIA) is worthy of further investigation. In this study, we further demonstrated that the cytotoxic concentration of ANE 30-100K induces some typical autophagy hallmarks in esophageal carcinoma (CE81T/VGH) cells in an Atg5-dependent manner. Furthermore, the endocytosis inhibitor (methyl-β-cyclodextrin) and two caveolin shRNAs, as well as two proteasome inhibitors (lactacystin and epoxomicin), were shown to attenuate ANE 30-100K-induced cytotoxicity and LC3-II accumulation significantly in OECM-1 and CE81T/VGH cells. Finally, we also analyzed the carbohydrate compositions of ANE 30-100K by phenol-sulfuric acid method and high performance anion exchange chromatography with pulse amperic detector. The results showed that ANE 30-100K contains about 67% carbohydrate and is composed of fucose (5.938%), arabinose (24.631%), glucosamine (8.066%), galactose (26.820%), glucose (21.388%), and mannose (13.157%). Collectively, these results suggest that caveolin-mediated endocytosis and proteasome are required for AIA and the major components of ANE 30-100K are carbohydrates. This study may have provided new knowledges of the action mechanisms and compositions of ANE 30-100K.

## Introduction

Betel quid (BQ) and areca nut (AN) are both human carcinogens; however, they are popular worldwide, especially in the Asia-Pacific regions. They are risk factors for both oral and esophageal cancers but unlike tobacco, no political or official restriction to these carcinogens has been documented in these countries. It is thought that further investigations of BQ- and AN-associated mechanisms may be required for the establishment of restriction policies for BQ and AN users [1].

AN extract (ANE) can induce the generation of reactive oxygen species (ROS) in both saliva and solution, as well as the modification of DNA [2, 3]. The major alkaloid of AN, arecoline, is capable of causing oral submucous fibrosis (OSF), oral squamous cell carcinoma (OSCC) and genotoxicity [4]. Moreover, arecoline and another AN ingredient, oligomeric procyanidins, are known to trigger cellular apoptosis [5, 6]. Unexpectedly, we noticed that after treatment of cells with ANE or its 30-100 kDa fraction (ANE 30-100K), the morphological changes including swollen cell, shrunken nucleus, and empty cytoplasm were distinct from those of apoptosis. ANE 30-100K induced the accumulation of microtubule-associated protein 1 light chain 3A/B-II (LC3-II), acidic vesicles, and autophagic vacuoles in OECM-1, indicating the induction of macroautophagy (hereafter referred to as autophagy) [7]. Similar responses to ANE 30-100K were also observed in oral fibroblasts, peripheral blood lymphocytes, and Jurkat T cells [8].

Roles of autophagy in tumor growth are important and complicated. In stromal cells, autophagy can provide nutrients to tumors; meanwhile, it helps tumors to survive intracellular and environmental stress. In contrast to these tumor-promotive roles, autophagy can remove abnormal mitochondria to prevent excessive ROS production and to maintain gene integrity, as well as support anticancer immune responses by regulating T cells and natural killer cells [9, 10]. These opposing roles of autophagy in tumor formation and progression might be probably the reason for the opposite outcomes of tumor therapy in mice through autophagy inhibition [11]. Thus, investigation of cellular responses to ANE 30-100K-induced autophagy (AIA) may achieve better understanding of AN-associated cancers.

Recently, we verified that AIA was clathrin- and dynamin-dependent [12], suggesting the initiation of AIA through a receptor- and clathrin-mediated endocytosis. In addition to clathrin-mediated endocytosis, another widely expressed and dynamin-dependent endocytosis is the caveolin-mediated endocytosis, regarded as an alternative route for the entrance of extracellular substances into cells [13, 14]. The fates of engulfed endosomes can be either fused with autophagosomes [15] or targeted to proteasome degradation as receptors of some cytokines and growth hormone upon ligand binding [16-18].

In this study, we monitored autophagic responses of the esophageal CE81T/VGH cells to ANE 30-100K, and the roles of caveolin-mediated endocytosis and proteasome in AIA were also analyzed. The effects of a endocytosis inhibitor (methyl-β-cyclodextrin, MβCD), two short hairpin RNAs (shRNAs) of caveolin 1, and two proteasome inhibitors (epoxomicin and lactacystin) on AIA were measured in OECM-1 and CE81T/VGH cells. In addition, the carbohydrate compositions of ANE 30-100K were also analyzed by phenol-sulfuric acid method and high performance anion exchange chromatography with pulse amperic detector.

## Materials and methods

### Chemicals

Methyl-β-cyclodextrin (128446-36-6), lactacystin (133343-34-7), and epoxomicin (134381-21-8) were purchased from Sigma-Aldrich (Saint Louis, MO, USA). XTT reagents (11465015001) were bought from Roche Molecular Biochemicals (Indianapolis, IN, USA).

### Preparation of ANE 30-100K

In Southeast Asian, mature (ripe) AN is commonly used for chewers; however, immature (tender) AN is also popular in Taiwan. We extracted one tender AN in a earthenware basin with 5 ml PBS (phosphate-buffered saline, containing 1.8 mM KH_2_PO_4_, 150 mM NaCl, 8.1 mM Na_2_HPO_4_, 2.7 mM KCl, pH 7.4) at room temperature, and the extracted solution was centrifuged at 12,000 *g* for 10 min at 4°C. The supernatant was used as ANE (areca-nut-extract). Next, ANE was filtered through cotton to remove the insoluble materials as much as possible and then centrifuged at 2,900 *g* for 30 min at 4°C with 100 kDa-pored concentration tube (Z677906, Sigma-Aldrich). The flow-through fraction was collected and further subjected to centrifugation as above with 30 kDa-pored concentration tube (430-30, Sigma-Aldrich). To remove small molecules like salts and phytochemicals, the materials of AN retained in 30 kDa-pored tube were washed twice with 2 ml H_2_O. The remaining substances were dissolved in water and lyophilized, designated as ANE 30-100K. Finally, the dried powder of ANE 30-100K was weighed and stored at −80°C.

### Cell culture and treatments

The esophageal carcinoma (CE81T/VGH) and o*ral* epidermoid carcinoma Meng-1 (OECM-1) cell lines were kindly and respectively provided by Dr. Cheng-Po Hu (Department of Medical Research and Education, Taipei Veterans General Hospital, Taipei, Taiwan) and Dr. Dar-Bin Shieh (Institute of Oral Medicine, National Cheng Kung University, Tainan, Taiwan), respectively. These two cells were cultured in Dulbecco’s modified Eagle’s medium (DMEM) (11995-065, Life Technologies Inc. Gibco/BRL Division, Grand Island, NY, USA) containing 10% fetal bovine serum (FBS) (16000-044, Life Technologies Inc.). Temperature of the incubator was set at 37°C in a humid atmosphere supplied with 5% CO_2_. Five thousand cells *per* well were cultured in a 96-well plate for cell viability determination with XTT {sodium 30-[1-(phenylaminocarbonyl)-3,4-tetrazolium]-bis (4-methoxy-6-nitro) benzene sulfonic acid hydrate} reagents by following manufacturer’s instruction.

For Western blotting, the 10-cm culture dish containing 5 × 10^6^ cells were used for lysate collection. Before any treatment, cells were serum starved overnight. The following treatment with chemicals (MβCD, epoxomicin, and lactacystin) and/or ANE 30-100K were also under serum-deprived conditions for 24 h. Cells were then scraped and treated with lysis buffer (0.5% NP-40, 2 mM EDTA, 50 mM Tris-HCl, pH 7.4) containing protease inhibitor cocktail set I (Calbiochem-Novabiochem Corp., San Diego, CA, USA). The collected lysate proteins were then subjected to immunoblotting with LC3 (L7543) and β-actin (A5441) antibodies (Sigma-Aldrich).

### Western blot analysis

Lysate proteins (20 μg) from treated cells were subjected to sodium dodecyl sulfate-polyacrylamide gel electrophoresis (SDS-PAGE) and then transferred onto a NC filter. This filter was then immunoblotted with the antibodies of LC3 (L7543, Sigma-Aldrich, 2000-fold diluted), Atg5 (2630, Cell Signaling Technology, Danvers, MA, USA, 1000-fold diluted), or β-actin (A5441, Sigma-Aldrich, 10,000-fold diluted). The secondary horseradish peroxidase-coupled goat anti-mouse-IgG (AP124P, Merck Millipore, Darmstadt, Germany) for β-actin antibody was 10,000-fold diluted and goat anti-rabbit-IgG (81-6120, Invitrogen Corporation, Camarillo, CA, USA) 2,000-fold diluted for LC3 antibody. Signals of LC3 and β-actin proteins were developed with Western Lightening Chemiluminescence Reagent (Perkin–Elmer Life Sciences, Inc., Boston, MA, USA) and then digitalized with UnSCAN-IT software Automated Digitizing System, version 5.1.

### Observation of morphological changes, acidic vesicles, and autophagic vacuoles

CE81T/VGH cells seeded in 96-well plate were treated with ANE 30-100K (10 μg/ml) for 0, 3, and 24 h. The morphological changes of these cells were firstly observed and photographed by light microscopy. We next tried to analyzed whether the intracellular vacuoles emerged after ANE 30-100K treatment for 3 h could be stained by acridine orange. Thus, these cells (seeded in slide chamber) were stained with acridine orange (1 μg/ml) for 10 min and photographed under a fluorescent microscope. The percentage of acidic vesicle-containing cells in randomly chosen 100 cells from three independent microscopic fields was determined. Finally, 3-h ANE 30-100K-treated cells were soaked in glutaraldehyde (2.5%) for 1 h and then in OsO_4_ (1%) for another 20 min at 4°C (fixation) and then dehydrated with a serial ethanol gradient [50% (10 min), 75% (10 min), 85% (10 min), and 95% (10 min, twice)], followed by propylene oxide (10 min, twice). These cells were impregnated with propylene oxide (50%) and embedding chemicals (Epon 812:Araldite 502:DDSA:DMP 30 = 1.34:0.93:2.60:0.14, Electron Microscopy Sciences, PA, USA) (50%) for 30 min, and then embedded in 100% embedding chemicals. Before examination by TEM (JEOL JEM-1200EX) at 80 kV, the ultrathin sections were stained with uranyl acetate and lead citrate and photographed with electron microscopy film 4489 170 Estar Thick Base (Kodak, NY, USA).

### Lentivirus-mediated shRNA interference

*E. coli* bacteria transfected with pLKO.1 plasmid containing shRNA sequences of human atg5 and caveolin 1 genes were bought from the National RNAi Core Facility (NRCF, Academia Sinica, Taipei, Taiwan). The target sequences of atg5 shRNA were CCTTTCATTCAGAAGCTGTTT (clone ID: TRCN0000151474; Region: CDS) and those of caveolin 1 were GCTTTGTGATTCAATCTGTAA (Clone ID: TRCN0000007999; Region: 3’UTR) and GACGTGGTCAAGATTGACTTT (Clone ID TRCN0000008002; Region: CDS). These insert sequences have been confirmed by Genomics Inc. (New Taipei City, Taiwan). Afterwards, the three constructed plasmids as well as empty pLKO.1 plasmid were sent to NRCF for package into lentivirus and the multiplicity of infection (MOI) was also determined. CE81T/VGH (5 × 10^6^ cells) were infected by plasmid-packaged virus (MOI = 4), polybrene (8 µg/ml) (107689, Sigma-Aldrich), and 1% FBS for 24 h. Cells were then selected in medium containing puromycin (2 µg/ml) and 10% FBS for 48 h.

### Carbohydrate compositional analysis

ANE 30-100K (3 mg) in 500 μl sample solution was analyzed by phenol-sulfuric acid method. It was mixed with 500 μl phenol solution and hydrolyzed by concentrated sulfuric acid (2.5 ml) and stirred for 10 sec. Finally, it was left at room temperature for 20 min and the absorbance at 490 nm was measured by spectrophotometer. The standard monosaccharide used were galactose and glucose.

Alternatively, polysaccharide compositional analysis (monosaccharide %) of ANE 30-100K (3 mg) was hydrolyzed with 4 N trifluoroacetic acid at 113°C for 12 hr and the residue was concentrated for compositional analysis by High Performance Anion Exchange Chromatography with Pulse Amperic Detector (HPAEC-PAD). ANE 30-100K (hydrolysates, 1 mg) was 100-fold diluted in double-distilled water and 10 μl of the sample was injected for HPAEC–PAD analysis. The above-prepared carbohydrate samples were analyzed using a DionexTM ICS-3000 DC equipment containing a gradient pump and an eluent degas module. Separation of carbohydrate molecules was carried out on a CarboPac PA-10 anion-exchange column (250 × 2 mm). The mobile phase contained 100 mM NaOH (eluent A) and 500 mM NaOAc (eluent B) in gradients. Eluent A was constant (100%) during 0–10 min and gradient (100% to 0%) was produced during 10–30 min with eluent B. The flow rate was 0.25 ml/min. Carbohydrates were detected by pulsed amperometric detection (PAD) with a gold working electrode and a hydrogen reference electrode. The temperature was set at 25°C and all analyses were carried out in duplicate.

### Statistical analysis

The OD_450_ values of XTT assay as well as LC3-II/β-actin and Atg5/β-actin ratios from three independent results between cells treated with ANE 30-100K alone and ANE 30-100K plus inhibitor (MβCD, epoxomicin, or lactacystin) were analyzed by one-way analysis of variance (ANOVA) with the Tukey Multiple Comparison Test (IBM SPSS Statistics 20.0.0 software). *P* value smaller than 0.05 was considered statistically significant.

## Result

### ANE 30-100K induces autophagic cell death of CE81T/VGH cells

ANE 30-100K has been shown to be cytotoxic to OECM-1 cells and the 50% inhibitory concentration (IC_50_) was 15 μg/ml [7]. Here, ANE 30-100K was also illustrated to reduce the viability of CE81T/VGH cells in a dose-dependent manner and the IC_50_ was 6.5 μg/ml (Fig 1A). Intriguingly, as observed by light microscope, some intracellular vacuoles emerged after ANE 30-100K treatment for 3 h and most of the cytoplasm became empty when the treatment extended to 24 h (Fig 1B). We next examined the intracellular vacuoles emerged at 3-h of ANE 30-100K treatment by acridine orange staining and transmission electron microscope (TEM). The results showed that after ANE 30-100K treatment for 3 h, intracellular vacuoles in CE81T/VGH cells stained by acridine orange exhibited orange-yellow color, i.e., the color of acidic vesicles (Fig 1C). Next, these cells when observed with TEM, showed typical double-membrane autophagic vacuole structures (Fig 1D). Some of these morphological changes are similar to those of OECM-1 cells as observed in our previous study, indicating that ANE 30-100K can stimulate a massive degradation of cytoplasmic materials in both OECM-1 and CE81T/VGH cells through autophagy [7].

**Fig 1.**
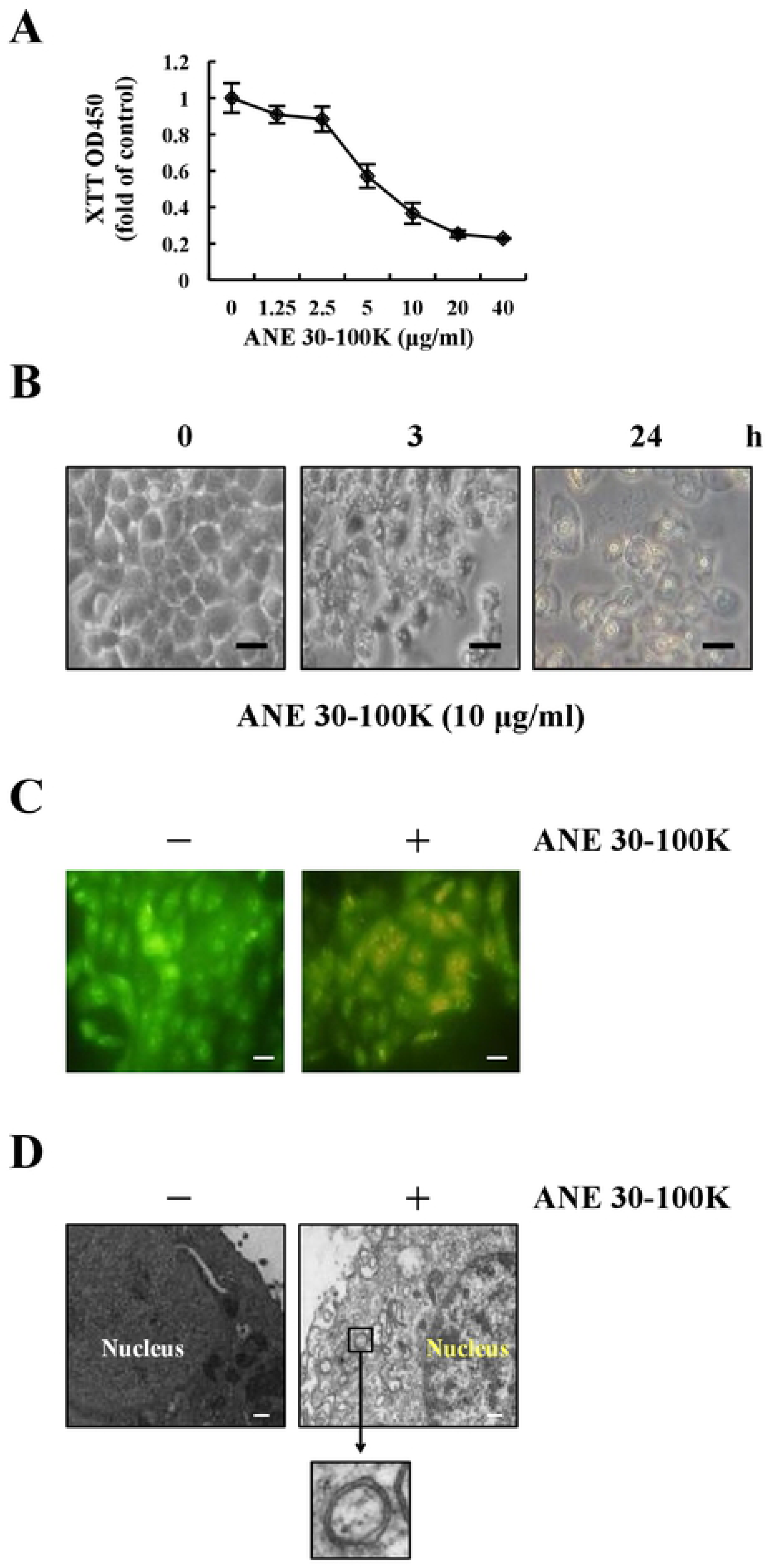
ANE 30-100K induces morphological changes of CE81T/VGH. (A) CE81T/VGH cells seeded in 96-well plate were serum starved overnight and treated with the indicated concentrations of ANE 30-100K for 24 h in a triplicated manner and subjected to XTT assay. Average OD_450_ absorbance of triplicated wells (± SD) were normalized against that of untreated control cells. (B) CE81T/VGH cells seeded in slide chamber were serum starved overnight and treated with ANE 30-100K (10 μg/ml) for the indicated periods. Cells were then observed and photographed under light microscope. Bar = 10 μm. (C) CE81T/VGH cells treated with or without ANE 30-100K (10 μg/ml) for 3 h were stained by acridine orange and photographed under a fluorescent microscope. Bar = 5 μm. (D) Alternatively, cells treated as in (C) were analyzed and photographed with a transmission electron microscope as described in materials and methods. One of the typical autophagic vacuoles with double-membrane structure was 16-fold further magnified and shown. Bar = 0.5 μm.

In addition to these observations, we also demonstrated that treatment of CE81T/VGH cells with ANE 30-100K for 24 h induced the elevation of LC3-II levels in both dose- and time-dependent manners (Fig 2A and 2B). Meanwhile, LC3-II level was further increased in the presence of lysosomal inhibitors including pepstatin A (10 μg/ml), E64d (10 μg/ml), and leupeptin (10 μg/ml) (Fig 2C). This feature is regarded as the induction of autophagic flux [19], and similar effect of ANE 30-100K on oral fibroblasts has also been demonstrated in our previous studies [8]. Finally, we also tried to utilize atg5 shRNA to inhibit the expression of Atg5 protein. Fig 3A showed that Atg5 protein level was significantly decreased in CE81T/VGH cells infected with atg5 shRNA-containing virus compared to non-infected cells (control) and cells infected with empty plasmid-containing virus (virus-plasmid control, VPC). Furthermore, these Atg5-knocked down cells became more resistant to cytotoxic ANE 30-100K challenge (for 24 h) as revealed by XTT assay (Fig 3B). The percentage of acidic vesicles was also significantly reduced in Atg5-knocked down cells than VPC and control cells after 3 h of ANE 30-100K stimulation (Fig 3C). Moreover, ANE 30-100K-induced LC3-II accumulation was nearly abolished in these Atg5-deficient cells (Fig 3D). Collectively, these results suggested that ANE 30-100K is capable of inducing autophagic cell death of CE81T/VGH in an Atg5-dependent manner.

**Fig 2.**
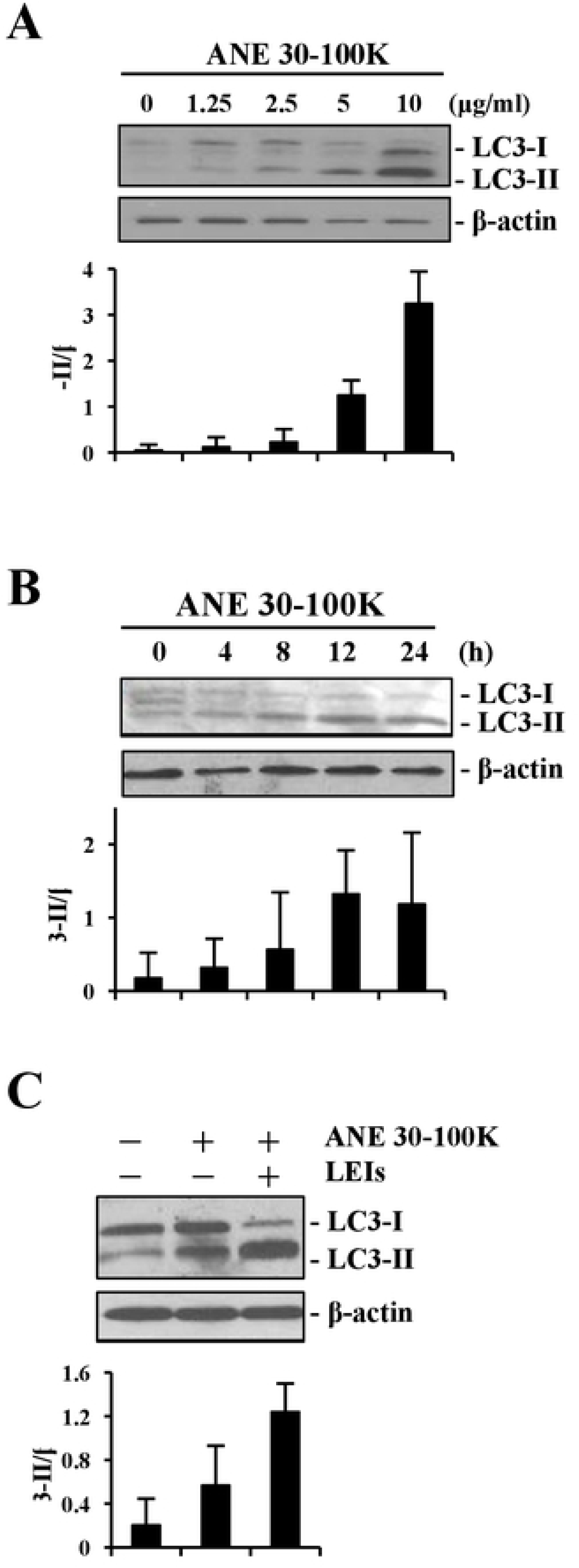
ANE 30-100K induces autophagic flux in CE81T/VGH. (A) CE81T/VGH cells treated with the indicated concentrations of ANE 30-100K as **Fig 1A** were subjected to immunoblotting with LC3 and β-actin antibodies. Protein signals from two independent experiments were digitalized and the average LC3-II/β-actin ratios ± SD were plotted under each lane. (B) CE81T/VGH cells treated with the indicated periods with ANE 30-100K (10 μg/ml) were subjected to the same immunoblotting and plotted as (A). (C) Untreated control CE81T/VGH cells and cells treated with ANE 30-100K (10 μg/ml) with or without lysosomal enzyme inhibitors (LEIs), containing pepstatin A (10 μg/ml), E64d (10 μg/ml), and leupeptin (10 μg/ml), were identically immunoblotted and plotted as (A).

**Fig 3.**
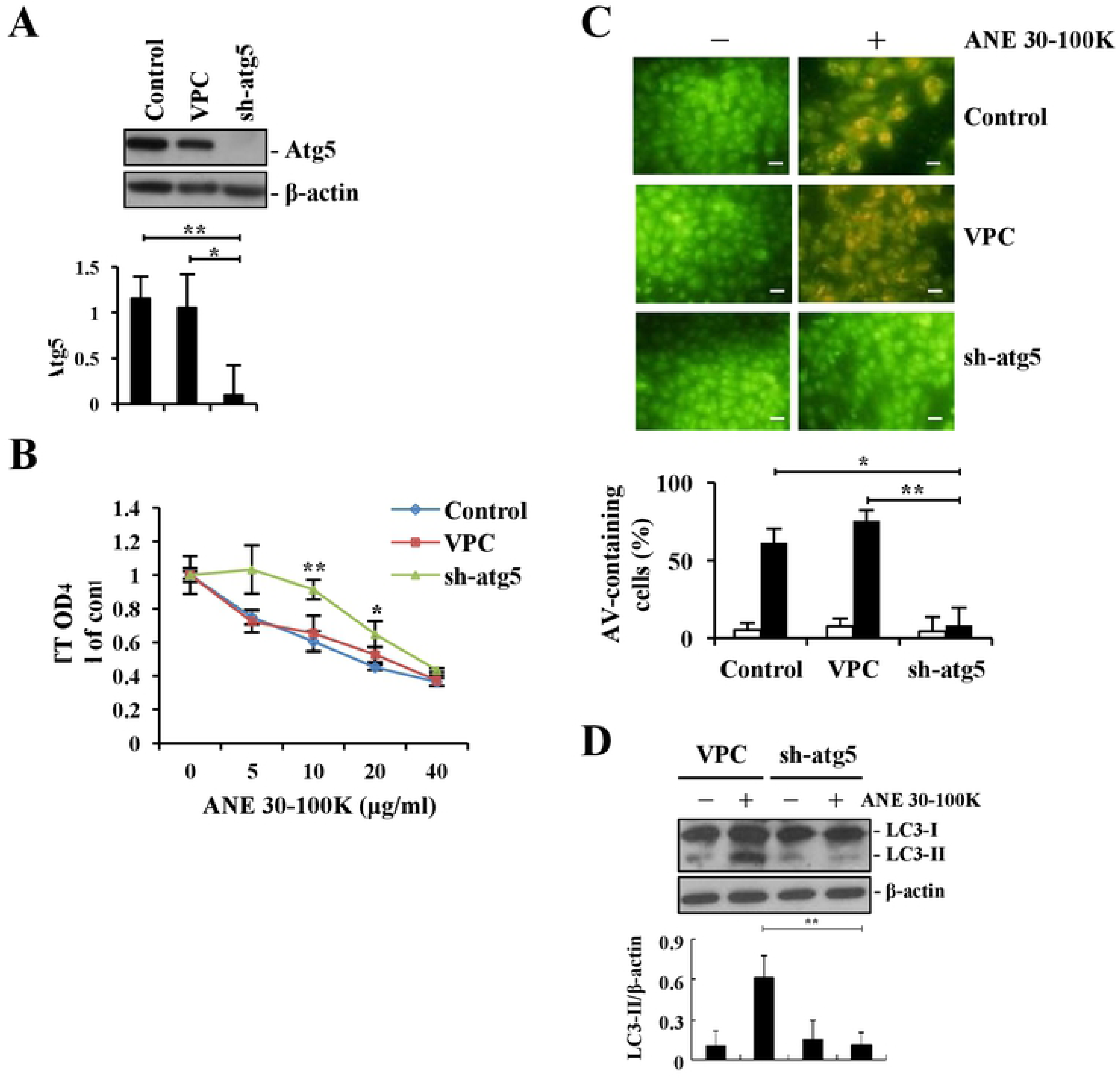
Knockdown of Atg5 expression in CE81T/VGH reduces ANE 30-100K-induced autophagic cell death. (A) Non-transduced control CE81T/VGH cells and cells infected with lentiviruses containing either empty pLKO.1-Puro plasmid (virus control, VC) or pLKO.1-Puro plasmid inserted with atg5 shRNA (sh-atg5) were subjected to puromycin selection. These cells were than immunoblotted with Atg5 and β-actin antibodies. Protein signals from two different independent immunoblots were digitalized and the average Atg5/β-actin ratios ± SD were plotted under each lane. (B) Control, VC, and sh-atg5 cells seeded in 96-well plate were treated with the indicated concentrations of ANE 30-100K for 24 h in a triplicated manner and analyzed by XTT reagents. (C) Control, VC, and sh-atg5 cells seeded in slide chamber treated with ANE 30-100K (10 μg/ml) for 3 h were stained by acridine orange and photographed under a fluorescent microscope. The acidic vesicle (AV)-containing cells in one hundred randomly chosen cells in a microscopic field were determined and the average percentage (± SD) from three different fields was obtained and plotted. Bar = 10 μm. (D) VC and sh-atg5 cells treated with ANE 30-100K (10 μg/ml) for 24 h were immunoblotted with LC3 and β-actin antibodies. Average ratios of LC3-II/β-actin from two independent experiments were plotted under each lane. *: *p* < 0.05, **: *p* < 0.01.

### MβCD, caveolin 1 shRNAs, lactacystin, and epoxomicin attenuate ANE 30-100K-induced autophagic cell death

MβCD can extract membrane cholesterol and is capable of inhibiting both clathrin- and caveolin-mediated endocytosis [20]. We firstly tested the cytotoxic effects of this compound on CE81T/VGH and OECM-1 cells, and found that up to 0.5 μM of MβCD, no significant cytotoxicity was observed in both cells (Fig 4A). Pretreatment of both cells with MβCD (0.5 μM) significantly reduced the cytotoxicity of ANE 30-100K (Fig 4B). Moreover, such pretreatment could also significantly attenuate ANE 30-100K-induced LC3-II accumulation (Fig 4C).

**Fig 4.**
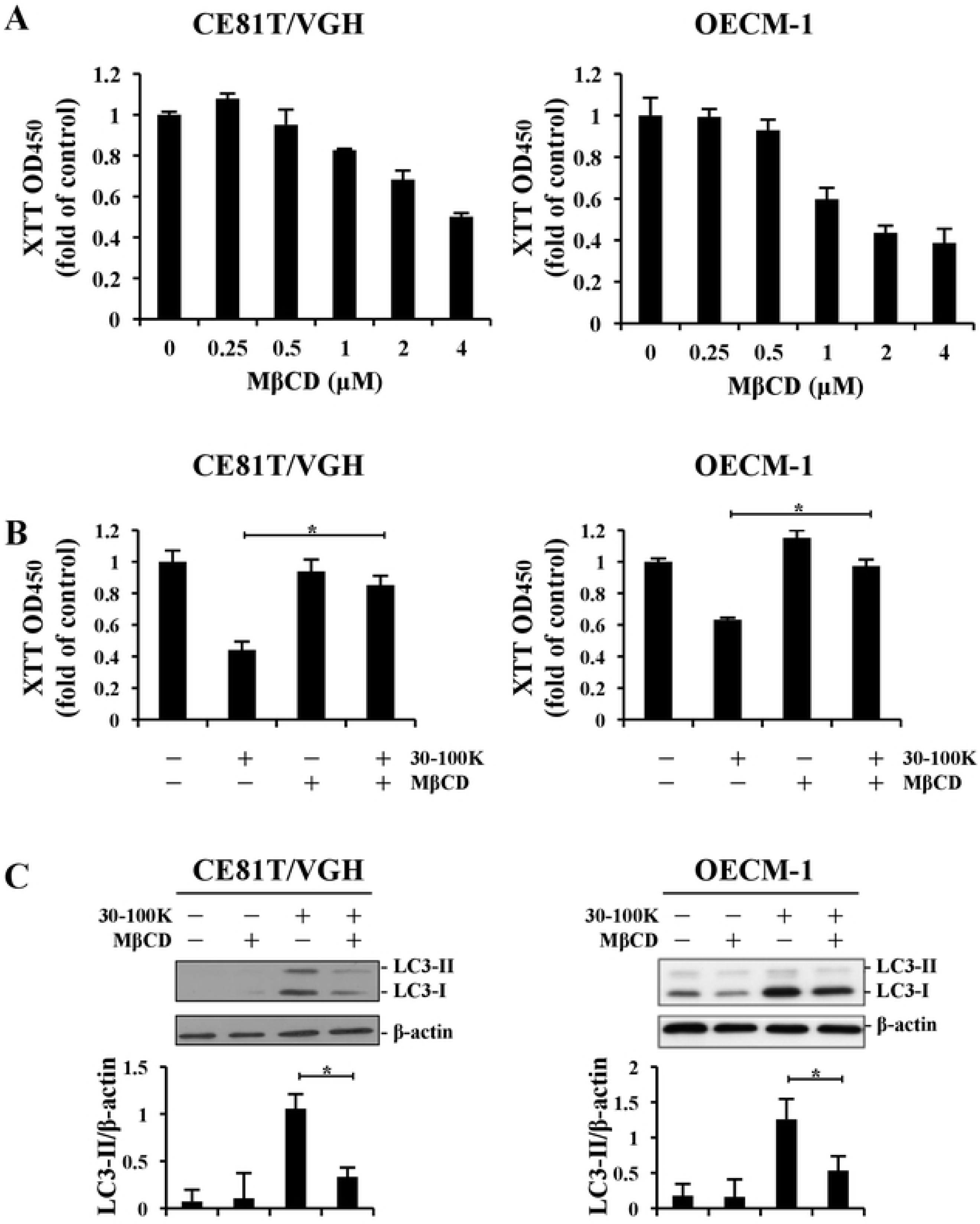
Methyl-β-cyclodextrin attenuates ANE 30-100K-induced autophagic cell death. (A) CE81T/VGH and OECM-1 cells treated with the indicated concentrations of methyl-β-cyclodextrin (MβCD) were analyzed by XTT assay. Average OD_450_ values from triplicated wells were plotted as mean ± SD. (B) CE81T/VGH and OECM-1 cells pretreated with MβCD (0.5 μM) were treated with or without ANE 30-100K (10 μg/ml) (30-100K) and analyzed as A. (C) CE81T/VGH and OECM-1 cells treated as in B were parallelly subjected to Western blot with LC3 and β-actin antibodies. Protein signals were digitalized and the average LC3-II/β-actin ratios (mean ± SD) were plotted under each treatment. *: *p* < 0.05.

Plasmid PLKO.1 was used to carry two different shRNAs targeting caveolin 1 cDNA 3’UTR (7999) and CDS (8002). Lentiviruses carrying empty plasmid (VPC), 7999 shRNA-inserted plasmid (7999), and 8002 shRNA-inserted plasmid (8002) were transduced into CE81T/VGH and OECM-1 cells. XTT analysis revealed that 7999- and 8002-infected CE81T/VGH and OECM-1 cells were more resistant to cytotoxic ANE 30-100K insult compared to uninfected cells (control) and VPC cells (Fig 5A). Moreover, ANE 30-100K-provoked LC3-II increase was also profoundly reduced in 7999-infected (Fig 5B) and 8002-infected (Fig 5C) OECM-1 and CE81T/VGH cells than in those of their respective control and VPC cells.

**Fig 5.**
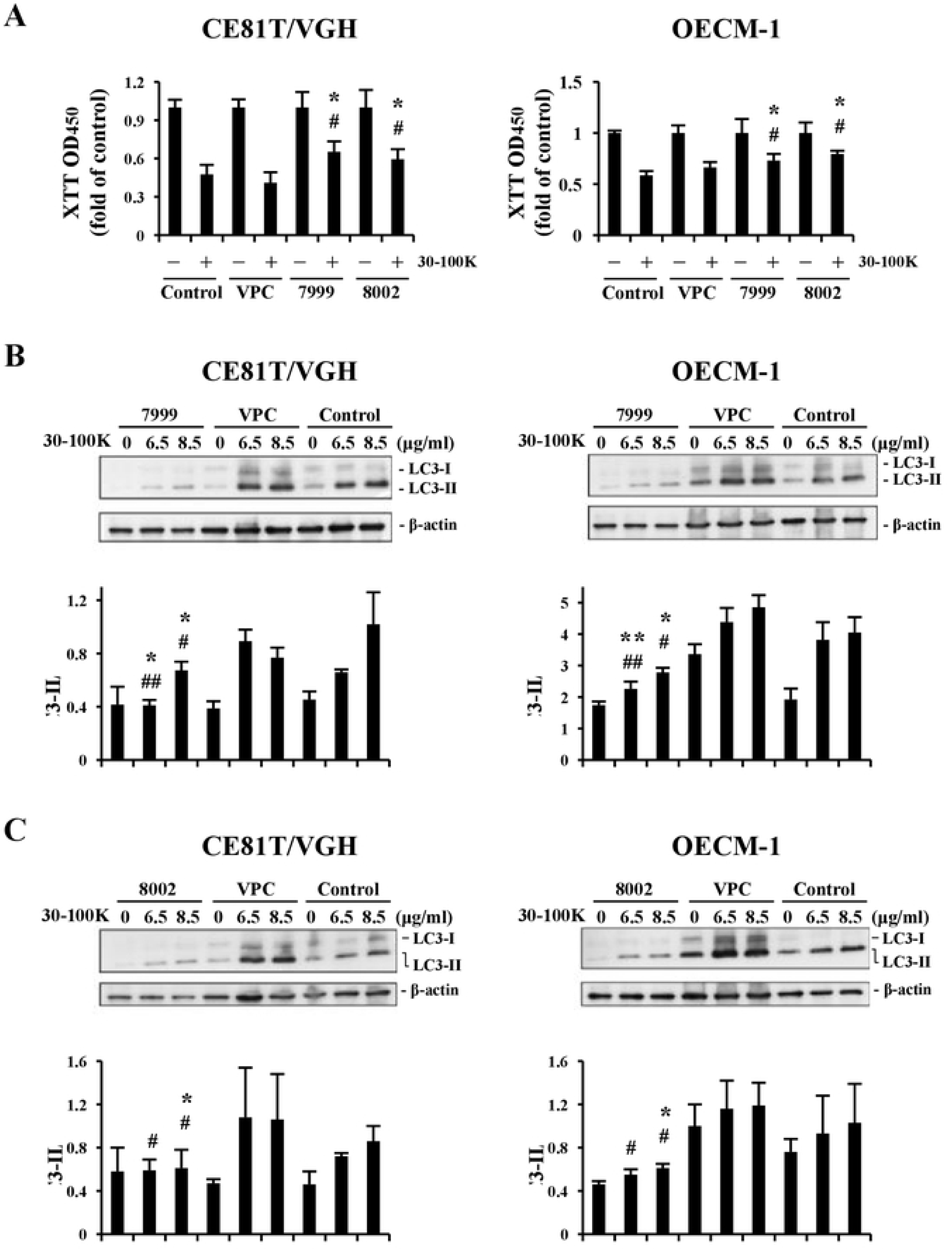
Caveolin 1 shRNA inhibits ANE 30-100K-induced autophagic cell death. (A) CE81T/VGH and OECM-1 cells (control) infected with empty plasmid (virus plasmid control, VPC) or plasmids containing caveolin 1 shRNA inserts 7999 and 8002 were treated with or without ANE 30-100K (10 μg/ml) (30-100K) and analyzed by XTT assay. Average OD_450_ values from triplicated wells were plotted as mean ± SD. (B) 7999, VPC, and control cells treated with the indicated concentrations of ANE 30-100K were subjected to Western blot analysis with LC3 and β-actin antibodies. Protein signals were digitalized and the average LC3-II/β-actin ratios (mean ± SD) were plotted under each treatment. (C) The results of 8002, VPC, and control cells subjected to the same treatment were presented as B. *: statistics vs. VPC, #: statistics vs. Ctr; * and #: *p* < 0.05, ** and ##: *p* < 0.01.

Lactacystin and epoxomicin are two selective proteasome inhibitors [21, 22]. We titrated their cytotoxic concentrations in CE81T/VGH and OECM-1 cells and found that no evident cytotoxicity were observed for lactacystin (2.5∼20 μM) in both cells; whereas 100 nM of epoxomicin began to show mild cytotoxicity to OECM-1 cells (Fig 6A). By either pretreating cells with lactacystin (5 μM) or epoxomicin (50 and 25 nM respectively for CE81T/VGH and OECM-1 cells) for 10 minutes, both cells exhibited stronger resistance against cytotoxic ANE 30-100K treatment (Fig 6B). In addition to cytotoxicity, we also measured the effects of lactacystin and epoxomicin pretreatment on ANE 30-100K-stimulated LC3-II accumulation. Fig 6C illustrates that pretreatment of both proteasome inhibitors significantly alleviated ANE 30-100K-induced LC3-II elevation in both cells.

**Fig 6.**
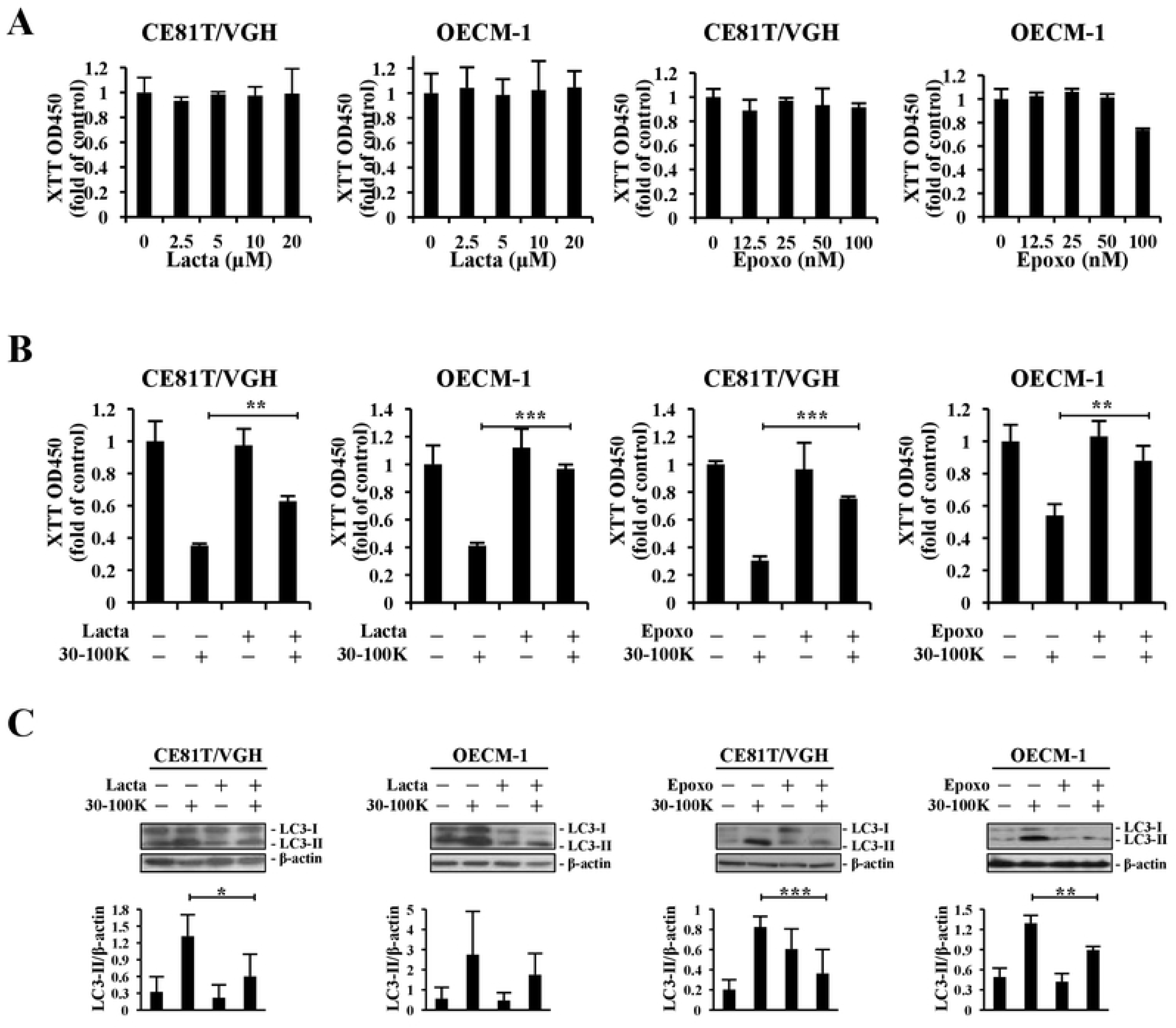
Lactacystin and epoxomicin reduce ANE 30-100K-induced autophagic cell death. (A) CE81T/VGH and OECM-1 cells treated with the indicated concentrations of lactacystin (Lacta) and epoxomicin (Epoxo) for 24 h were subjected to XTT assay. Average OD_450_ values from triplicated wells were plotted as mean ± SD. (B) These two cells treated with Lacta (5 μM) for 10 min were incubated with ANE 30-100K (10 μg/ml) (30-100K) for 24 h and then subjected to XTT assay. On the other hand, CE81T/VGH and OECM-1 cells respectively pretreated with 50 and 25 nM of Epoxo for 10 min, followed by the same 30-100K incubation were also analyzed by XTT assay. Average OD_450_ values ± SD were plotted as A. (C) Lysate proteins from CE81T/VGH and OECM-1 cells identically treated as B were subjected to Western blot analysis with LC3 and β-actin antibodies. Protein signals were digitalized and the average LC3-II/β-actin ratios (mean ± SD) were plotted under each treatment. **: *p* < 0.01, ***: *p* < 0.001.

### Carbohydrate composition of ANE 30-100K

Analysis using pheno-sulfuric acid method showed that ANE 30-100K contained 66% and 67% carbohydrate when galactose and glucose were used as the standard monosaccharide, respectively. Analytical results of HPAEC-PAD was illustrated in Fig 7., peak 1=fucose (5.938%), peak 2=arabinose (24.631%), peak 3=glucosamine (8.066%), peak 4=galactose (26.820%), peak 5=glucose (21.388%), peak 6=mannose (13.157%).

**Fig 7.**
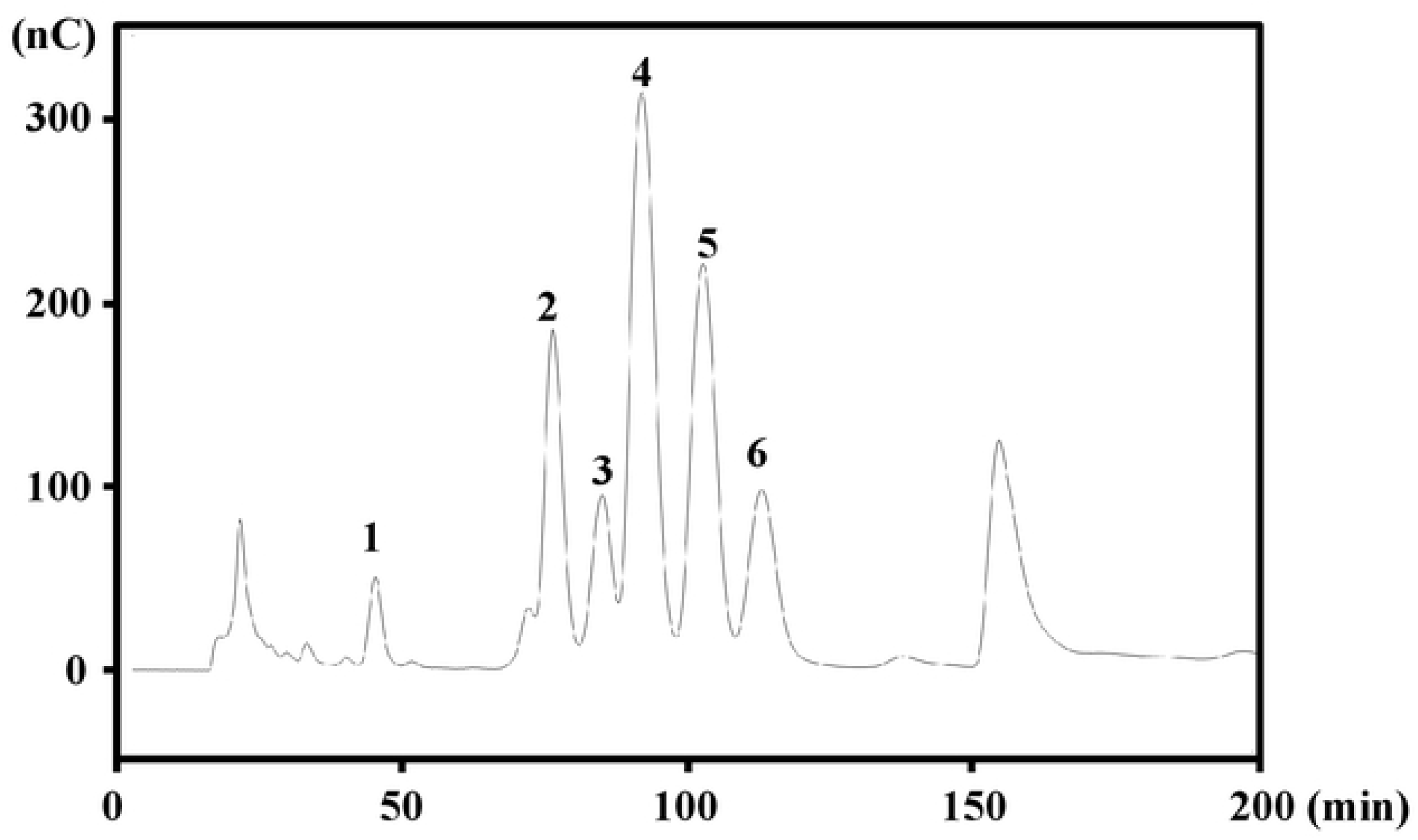
ANE 30-100K contains fucose, arabinose, glucosamine, galactose, glucose, and mannose. Monosaccharide composition of ANE 30-100K (3 mg) was analyzed by HPAEC-PAD as described in Materials and methods. Peaks 1-6 are fucose (5.938%), arabinose (24.631%), glucosamine (8.066%), galactose (26.820%), glucose (21.388%), and mannose (13.157%), respectively.

## Discussion

Together with our previous studies, we have further confirmed that ANE 30-100K not only induces autophagic cell death in OECM-1 cells, oral fibroblasts, and lymphocytes [7, 8], but also in CE81T/VGH (Fig 1∼Fig 3). It is thus thought that ANE 30-100K is capable of inducing autophagy in a wide range of different cell types. Moreover, we recently demonstrated that AIA is sensitive to benzyl alcohol (a membrane fluidity blocker), dynasore (a specific dynamin inhibitor), and shRNAs of dynamin and clathrin [12]. Here, we further show that AIA is also sensitive to MβCD and caveolin 1 shRNAs. Collectively, these data indicated that the autophagy-inducing AN ingredient (AIAI) in ANE 30-100K may enter cells through cholesterol-, caveolin-, dynamin-, and clathrin-dependent endocytosis and trigger the following autophagy program. Since these endocytic machineries are widely expressed in different types of cells, it is expected that most, if not all, types of cells can uptake AIAI and initiate autophagy responses.

It is reasonable to speculate the engulfment of AIAI through receptor-mediated endocytosis. However, whether there is a specific AIAI receptor on cell membrane remains unknown. Some membrane receptors have been reported to initiate autophagy. For examples, toll-like receptor (TLR)-mediated signals in macrophages has been linked the autophagy pathway to phagocytosis [23]. Moreover, inactivated epidermal growth factor receptor (EGFR) is known to induce autophagy, and Tan *et al.* recently showed that the oncoprotein LAPTM4B (lysosomal-associated protein transmembrane-4β) promoted a kinase-independent role for EGFR in autophagy initiation [24]. Pan *et al.* also reported that capsicodendrin, a natural compound isolated from *Cinnamosma macrocarpa*, strongly induced autophagy directly through inactivation of VEGFR2 (vascular endothelial growth factor receptor 2) and downstream AKT signaling [25]. Thus, it is intriguing to address whether AIAI binds to TLR or inactivates EGFR and/or VEGFR2 to trigger autophagy. Still, it is also possible for AIAI to interact with some other unknown membrane proteins to turn on autophagy program.

In fact, the autophagy-inducing activity of AIAI has been verified to be sensitive to lysing enzymes (including cellulase, β-glucanase, chitinase, and protease), cellulase, and proteinase K digestion in our previous study, suggesting that AIAI contains both protein and carbohydrate moiety, i.e., a proteoglycan or glycoprotein. After analyzing ANE 30-100K with Molish’s Test, Seliwanoff’s Test, thin-layer chromatography, and Protein Assay Kit (Bio-Rad Laboratories), it was shown to be mainly composed of carbohydrates rather than proteins [26]. Here, we further revealed that ANE 30-100K contains about 67% carbohydrate (Fig 7). These results collectively suggest a possibility of AIAI to be a proteoglycan. Interestingly enough, a soluble proteoglycan, decorin, was recently shown to upregulate paternally expressed gene 3 and evoke autophagy in endothelial cells by direct interaction with VEGFR2, a process thought to be intrinsically involved in cancer initiation and progression [27]. Therefore, these results raised the possibility that AIAI might induce autophagy through mechanisms similar to decorin.

In Fig 6, effective attenuation of ANE 30-100K-induced autophagic cell death by epoxomicin and lactacystin implied the involvement of the proteasome in AIA. Epoxomicin (50 and 25 nM for CE81T/VGH and OECM-1, respectively) was more effective than lactacystin (5 μM for both cells) in blocking AIA. This could be due to efficient inhibitory effect of epoxomicin on chymotrypsin-like activity of proteasome, demonstrated to be 80-fold faster than lactacystin [28]. These proteasome inhibitors have been illustrated to impair the internalization of IL-2/IL-2 receptor (IL-2R) complex and prevent the lysosomal degradation of IL-2. It was concluded that proteasome is required for optimal endocytosis of the IL-2/IL-2R and the subsequent lysosomal degradation of IL-2, possibly by blocking trafficking to the lysosome [17]. Furthermore, inhibition of proteasome also blocked glutamate agonist AMPA-induced internalization of glutamate receptors [18]. Therefore, epoxomicin and lactacystin might have the possibility to prevent internalization of AIAI receptors to impede AIA.

In conclusion, we present new evidences here to reveal the requirement of caveolin and proteasome in the pathway of AIA. Moreover, the carbohydrate moiety of AIAI may be composed of fucose, arabinose, glucosamine, galactose, glucose, and/or mannose.

## Acknowledgements

We thank Dr. Wen-Bin Yang (Genomics Research Center, Academia Sinica) for analyzing the carbohydrate composition of ANE 30-100K.

